# Amplicon-guided isolation and cultivation of previously uncultured microbial species from activated sludge

**DOI:** 10.1101/2023.07.13.548900

**Authors:** Maarten D. Verhoeven, Per H. Nielsen, Morten K. D. Dueholm

**Affiliations:** Center for Microbial Communities, Department of Chemistry and Bioscience, Aalborg University, Aalborg, Denmark

## Abstract

Microbes are fundamental for biological wastewater treatment. However, most microbial species found in activated sludge (AS) from wastewater treatment plants (WWTPs) have never been isolated and grown as pure cultures, thus limiting our understanding of the underlying biological processes. To change this, we here introduce an experimental setup where the plating of dispersed AS bacteria are combined with 16S rRNA gene amplicon sequencing of total plate biomass for rapid identification of growth conditions that allow for the isolation of key microbial species in AS. We show that agarose plates composed of AS fluid supplemented with various carbon sources support the growth of many previously uncultivated AS bacteria. To confirm that the approach can also be used to isolate previously uncultured species, we picked 200 colonies from the plates for growth in liquid medium. This resulted in 185 growing cultures representing 102 strains based on unique 16S rRNA gene V1-V3 amplicon sequence variants (ASVs). Classification of the ASVs with the MiDAS 4 database revealed 48 distinct genera, including the previously uncultured AAP99, *Ca*. Propionivibrio, Ellin6067, midas_g_12, and *Ca*. Brachybacter. Among the ASVs that obtained species-level classification, we observed 43 unique species of which 29 were only classified based on the MiDAS placeholder taxonomy highlighting the potential for culturing many novel taxa. Preparation of glycerol stocks and subsequent validation by restreaking on plates resulted in 10 pure cultures of which six represent core or conditional rare or abundant (CRAT) species observed within the MiDAS global survey of WWTPs.

**Importance:** Biological wastewater treatment relies on complex microbial communities that assimilate nutrients and break down pollutants in the wastewater. Knowledge about the physiology and metabolism of bacteria in wastewater treatment plants (WWTPs) may therefore be used to improve the efficacy and economy of wastewater treatment. Our current knowledge is largely based on 16S rRNA gene amplicon profiling, fluorescence in situ hybridization studies, and predictions based on metagenome-assembled genomes. Bacterial isolates are often required to validate genome-based predictions as they allow researchers to analyze a specific species without interference from other bacteria and with simple bulk measurements. Unfortunately, there are currently very few pure cultures of microbes commonly found in WWTPs. To address this, we introduce an isolation strategy that takes advantage of state-of-the-art microbial profiling techniques to uncover suitable growth conditions for key WWTP microbes. We furthermore demonstrate that this information can be used to isolate key organisms representing global WWTPs.

## Introduction

Wastewater treatment is a vital technology in urbanized areas, as it protects public health, the environment, and enables resource recovery. The most common wastewater treatment process worldwide is the conventional activated sludge (AS) process. This process relies on complex microbial communities that grow as suspended aggregates using nutrients from the wastewater as feed and converting it into excess biomass. This biomass is then separated from the cleaned effluent through sedimentation (1). By understanding the physiology and metabolic potential of the microbes common in wastewater treatment plants (WWTPs), we can improve the efficacy and stability of wastewater treatment, reduce the release of greenhouse gases, and recover valuable resources like nitrogen, phosphorus, and biopolymers (1–3).

The emergence of widely accessible DNA sequencing technologies have dramatically improved our knowledge on the distribution, diversity and phylogeny of microorganisms present in different environments on Earth (4). Two major studies have investigated the global diversity of bacteria in WWTPs, the Global Water Microbiome Consortium (GWMC) project (5), and the Global Microbial Database for Activated Sludge (MiDAS) project (6). The latter features a full-length 16S rRNA gene amplicon sequence variant (FL-ASVs) resolved reference database for all common bacteria and archaea in WWTPs worldwide (MiDAS 4). This database contains a unique seven-rank (domain to species) taxonomy for all reference sequences with reproducible placeholder names for environmental taxa lacking official taxonomic classifications (6, 7). Using this database, the abundance of all community members in WWTPs, as well as information about their phylogenetic relation to known species has been described (6). However, our current understanding of their metabolism and community function is still limited, as it is based only on a small subset of species for which isolates, complete genomes, and physiological data are available (8).

Although advances in obtaining metagenome-assembled genomes (MAGs) have shed light on the ecological roles of some uncultured microbial species, as exemplified by the recent recovery of more than a thousand high-quality (HQ) MAGs from Danish WWTPs (9), predicting their physiology becomes increasingly challenging as more novel and distinct taxa are discovered. This is because many of the protein-encoding sequences in these novel microbes show low similarity with those that have been experimentally characterized in previous research. To further advance our understanding of individual bacteria in WWTPs, it is crucial to conduct more wet lab studies that can verify genomics-based predictions. This is especially important for species for which data on their physiology, metabolism, and cell biology is not currently available (4).

While metabolomics, proteomics, transcriptomics, and in-depth physiological studies of mixed or enriched cultures are possible, the results are often challenging to interpret as the community dynamics complicate pinpointing the specific traits of the species of interest. Moreover, it is not possible to stock and reproduce active biomass from environmental samples, which is required to ensure experiments can be reproduced. It is therefore import that we obtain pure cultures from the species of interest in AS (1, 4, 10).

At present, the vast majority of the bacterial species that make up the microbial communities in AS have not been cultured individually, and as such very few representative pure cultures are available (11). Specifically, we would like to isolate representatives for the core and conditional rare or abundant taxa (CRAT) found in WWTPs across the globe, as these are assumed to have the highest impact on the treatment efficacy (6). While there are a few studies available in which AS core bacterial species, such as *Acidovorax caeni* (12)*, Microthrix parvicella* (13)*, Nitrospira defluvii* (14), and *Zoogloea caeni* (15) were isolated, in most cases available AS isolates represent low abundant species in situ, and may therefore not contribute significantly to the microbial community in WWTPs.

The discrepancy between the number of species that can be cultured in the lab compared to the vastly higher number being present in the samples from a natural habitat has been known for decades and often is referred to as “the great plate anomaly” (4, 16). The difficulty of isolating species can be attributed to two primary factors: (i) the challenges associated with dispersing bacterial aggregates and separating individual cells, and (ii) the difficulty in predicting and recreating the specific environmental conditions necessary for the proliferation of targeted microbial community members (17).

In the present study, we address these challenges by introducing a simple isolation strategy where AS bacteria dispersed using sonication, filtered to remove aggregates, and subsequently plated on growth media based on AS fluid (ASF) supplemented with various carbon sources are combined with 16S rRNA gene amplicon sequencing of total plate biomass for rapid identification of growth conditions that allow for the isolation of individual microbial community members. Tailoring the medium composition closely toward the conditions experienced in WWTPs resulted in a higher species isolation yield compared to traditional media, allowing for the isolation of previously uncultured bacteria.

## Results and Discussion

### Preparation of single cell suspensions from activated sludge flocs

Most microbes within AS grow in flocs with multiple species. Cultivation of single strains, therefore, requires dispersal of the flocs, preferably without significant impact on the overall viability of bacterial cells present. To disrupt the flocs, AS samples obtained from the Aalborg West (AAW) WWTP were homogenized and sonicated, after which each sample was passed sequentially through 40 µm and 5 µm cell strainers. LIVE/DEAD staining of the dispersed samples showed predominantly single cells, and there was no detectable difference in viability between the single cell suspension compared to the source AS (Figure 1a). Furthermore, 16S rRNA gene amplicon sequencing on DNA extracted from AS and derived single cell suspension did not reveal any loss of species (ASV richness) during the preparation process, and the overall diversity was actually slightly higher in the single cell preparation (Figure 1b and Figure S1a). Community composition analysis showed that specific genera, such as *Ca*. Phosphoribacter (*Tetrasphaera* midas_s_5 in the MiDAS 4.8.1 taxonomy), *Ca*. Microthrix, and *Nitrospira* decreased in relative abundance across all replicates, whereas others, such as *Rhodoferax*, Ca. Brachybacter (OLB8 midas_s_29 in the MiDAS 4.8.1 taxonomy), midas_g_171, and *Acidovorax* increased (Figure 1c and Figure S1b). The decreased relative abundance of highly abundant AS genera explains the increased diversity observed for the single cell preparations.

**Figure 1.**
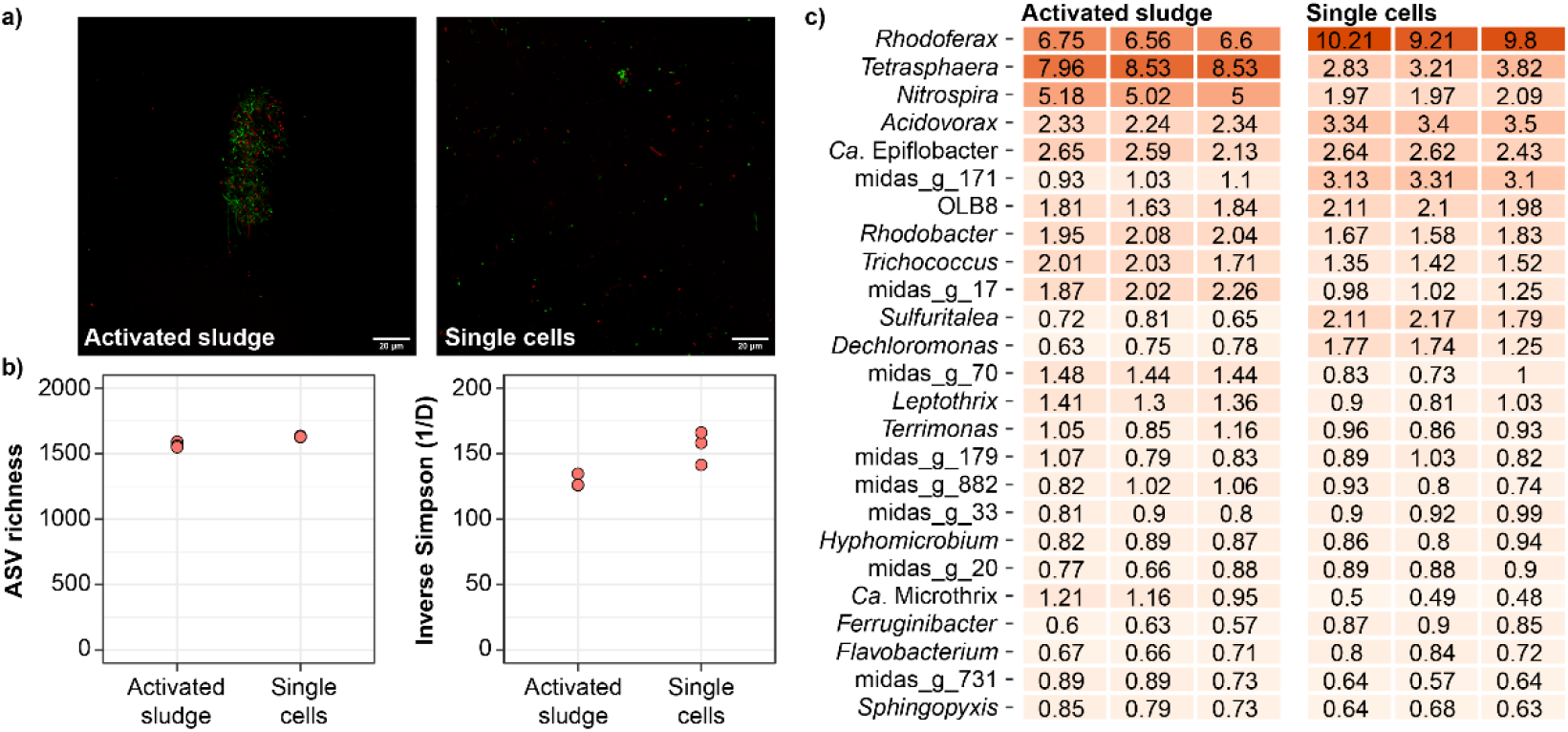
Preparation of single cell suspensions from activated sludge. a) Microscopy images of LIVE/DEAD-stained untreated activated sludge and single cell suspension. Live cells appear in green and dead in red. b) Alpha diversity based on 16S rRNA gene V1-V3 amplicon data for technical triplicates. c) Heatmap of the 25 most abundant genera in the activated sludge and single cell suspensions based on the V1-V3 amplicon data. All figures are based on activated sludge collected the 16th of December 2020 and used for oxic cultivations.

### AS fluid support the growth of previously uncultured species

To mimic the environmental conditions found in the WWTP, we extracted ASF directly from the source AS. For this, the supernatant of settled AS from the AAW WWTP was ultra-filtrated to yield a clear solution without particulate matter. Subsequently, the filtrate was concentrated by reverse osmosis and the retentate filter sterilized. The resulting ASF was supplemented to agarose culture plates with a variety of carbon sources in low concentration to mimic the oligotrophic environment encountered by the AS bacteria in situ (18). Agarose was used to eliminate any potential effects of impurities present in more regularly used agar (19, 20). Ammonium was added as an additional nitrogen source, except for plates with tryptone or casamino acids. Approximately 1000 single cells derived from AS were spread on each plate and these were incubated for 2-3 weeks under either oxic or anoxic conditions at 25°C.

To determine the effect of the different substrates, we performed 16S rRNA gene amplicon sequencing on DNA extracted from the entire biomass scraped off each agarose plate. Taxonomic classification of the resulting ASVs with the MIDAS 4.8.1 reference database allowed for genus- and species-level classification for most ASVs observed. Supplementing conventional culture media (R2A and TSB), which have previously been used for growing and isolating bacteria from activated sludge (10, 21), with ASF had a strong positive effect on the ASV richness and diversity (Figure 2a). The effect could also be observed directly on the agarose plates based on the diversity of colony morphologies (Figure S2). Compared to the conventional bacterial culture media, the ASV diversity that emerged on ASF plates supplemented with single carbon sources was substantially higher (Figure 2a).

**Figure 2.**
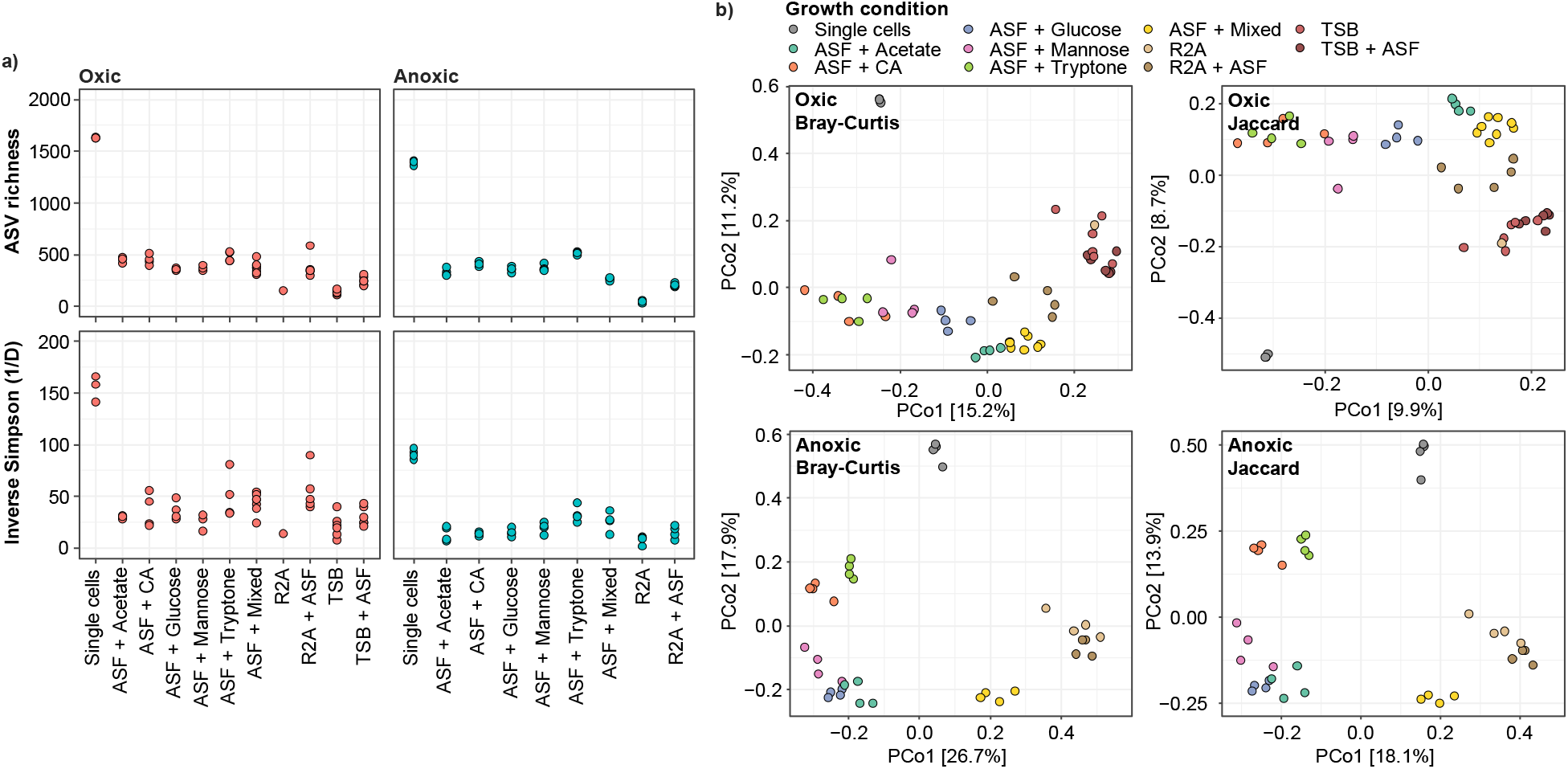
Microbial diversity of activated sludge single cell suspensions grown on various agarose media under oxic or anoxic conditions. a) Alpha diversity based on 16S rRNA gene V1-V3 amplicon data. b) Beta diversity based on 16S rRNA gene V1-V3 amplicon data. Bray-Curtis and Jaccard diversity were calculated at the ASV-level. ASF: Activated sludge fluid; CA: Casamino acids.

To further investigate the effect of media composition on the growth of the AS bacteria, we performed beta diversity analyzes (Figure 2b). PCoA plots of Bray-Curtis and Jaccard distances for ASVs revealed that each growth condition promoted the growth of a specific subset of the AS bacteria, and that similar carbon sources such as tryptone and casamino acids, or glucose and mannose, resembled each other more closely when it comes to ASV diversity and abundance. This suggests that specific growth conditions should be considered for targeted isolation of specific species. The sample clustering was more pronounced under anoxic compared to oxic conditions, which suggest that fermentative bacteria in general are more substrate specific.

To pinpoint which media promotes the growth of abundant AS bacteria, we compared the relative abundance of the most abundant genera in the dispersed AS with those obtained from the agarose plates (Figure 3). Because we only spread approx. 1000 bacterial cells on each agarose plate, and the harvested microbial biomass was visible by eye, we assume that any detectable reads in the amplicon data relates to actual growth of the associated species. With this in mind, we were able to detect growth of up to 64 of the top 100 most abundant genera in the dispersed AS using medium with the ASF, whereas media without ASF only allowed growth of up to 18 genera. This clearly demonstrates the importance of mimicking the source environment when trying to isolate new bacteria from a specific ecosystem.

**Figure 3.**
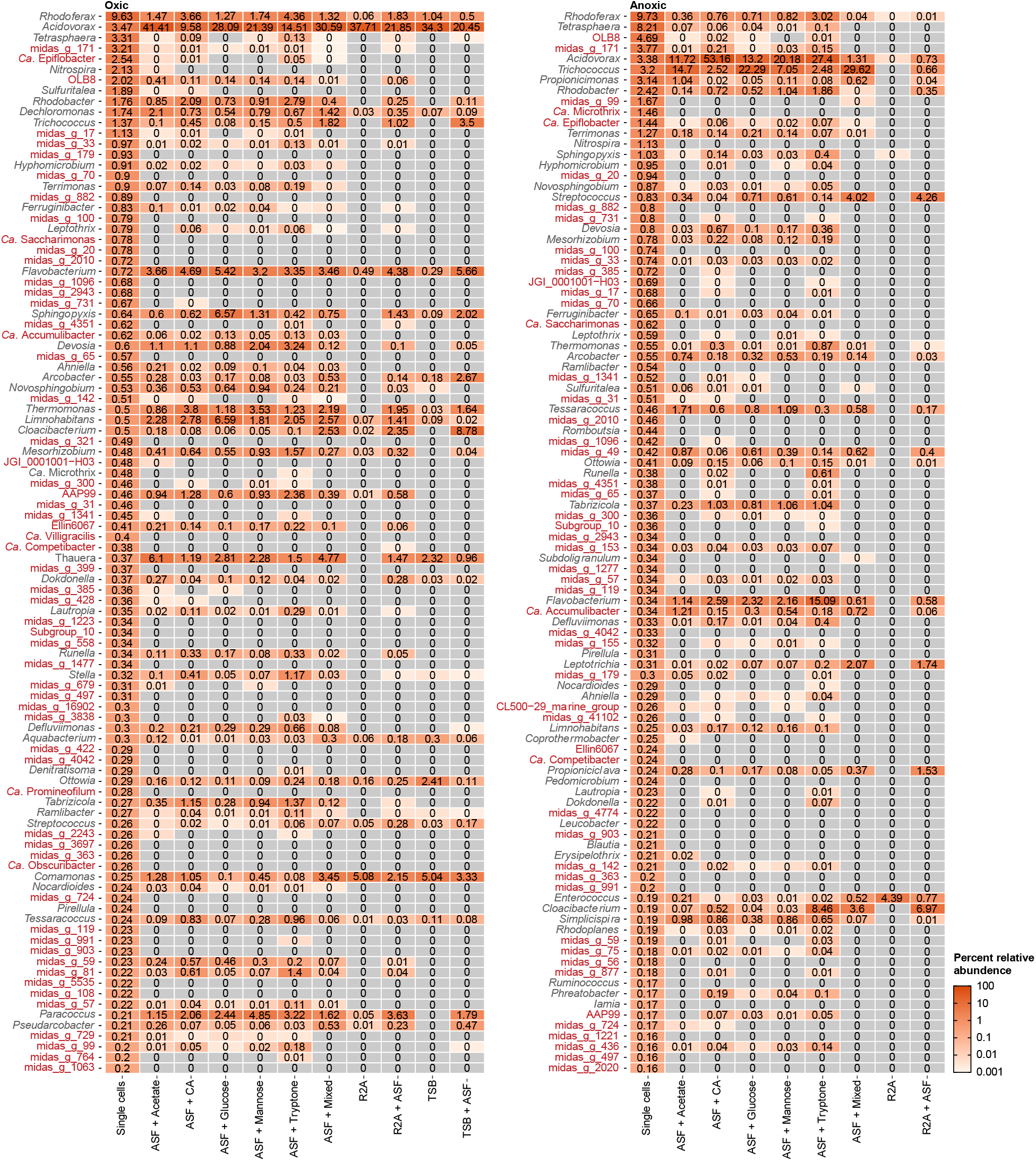
Heat map showing the top 100 genera found in the AS and the corresponding abundancy values found on each plate incubation under oxic and anoxic conditions. Values are the average abundance from four separately processed agarose plates. Genera colored in red have no isolated representatives. ASF: Activated sludge fluid; CA: Casamino acids.

Interestingly, ASF was able to stimulate the growth of several candidate genera and genera only described based on MiDAS placeholder taxonomy (Figure 3). These include genera of key importance for the wastewater treatment process, such as the polyphosphate accumulating organisms (PAOs) *Ca*. Accumulibacter and *Ca*. Phosphoribacter (*Tetrasphaera* midas_s_5 in the MiDAS 4.8.1 taxonomy). Because these genera have been key targets for isolation for many years, we investigated these genera in depth at the ASV-level (Figure S3). Unfortunately, we found that most of the ASVs classified as *Ca*. Accumulibacter were associated with the MiDAS placeholder species midas_s_168 and midas_s_3472, which were recently shown not to be PAOs and affiliated to a closely related genus that was named *Ca*. Propionivibrio (22). The most common *Ca*. Accumulibacter species, *Ca*. A. phosphatis was not able to grow under any of the applied conditions although present in the dispersed AS. It is, therefore, expected that this species lacks a vital nutrient in the media or a syntrophic partner. For *Tetrasphaera,* we observed growth of the midas_s_5, which represent the newly discovered *Ca*. Phosphoribacter, which is the most abundant PAO worldwide (23). In addition, we observed growth of midas_s_45 which belongs to the recently described *Ca*. Lutibacillus genus and represents another common PAO (23). Accordingly, it should be possible to isolate members of these genera.

The discrepancy between AS abundancy and the low ability of *Ca.* Phosphoribacter to grow on culture plates could be due to several factors. While 16S rRNA gene amplicon sequencing of the single cell material confirmed the presence of midas_s_5, it cannot be ruled out that the sonication treatment of the AS has disproportionately impacted the viability of this species (24). Detailed investigation on the impact of sonication on species specific viability could provide insight into how to obtain live single cells from the bacteria of interest.

Visual inspection of the culture plates prior to scraping colonies showed substantial variation in colony size (Supplemental Figure S2). Since the 16S rRNA gene amplicon data solely provides the relative abundance for growth on plates, it underrepresents slow growing organisms forming small colonies. Long term survey data from Danish WWTPs shows that midas_s_5 is highly enriched (up to 30%) within some plants while only present at low levels in sewage influent (0.05%). Nonetheless, 16S rRNA gene amplicon sequencing data (Figure 3) implies that midas_s_5 is likely forming colonies, albeit likely of small size.

### Competition from fast growing bacteria

Although many AS bacteria were able to grow on the ASF based agarose plates, their relative abundance on the plates was in general low. To learn more about the competing bacteria, we investigated the genera with the highest relative abundance on the agarose plates (Figure S4). These were *Acidovorax*, *Pseudomonas, Diaphorobacter,* and *Flavobacterium* under oxic conditions, and *Lactococcus* (mainly on R2A), *Acidovorax*, *Uliginosibacterium,* and *Trichococcus* under anoxic conditions (Figure 3).

Overall, it seems that the cultivation conditions on plates with ASF stimulated growth of many organisms that are more abundant in sewage compared to AS (25, 26). This indicates that the growth conditions on plates more closely resemble a nutrient rich sewer system environment than the oligotrophic environment found in the WWTPs, where the latter inherently select for microbes that manifest a kinetics-based strategy for proliferation (k-strategists) (18). Accordingly, culturing on plates is more suited for isolating rate-based strategists (r-strategists), since, with readily available substrate and virtually no competition from other cells, colonies can grow unhindered (4). *Acidovorax* has been commonly observed in WWTP that manifest a short sludge retention time (27). Moreover, it is one of the most abundant genera in Danish influent wastewater (25, 26). Yet, in most Danish WWTPs it seems that *Acidovorax* is not proliferating in the AS system and their presence is likely largely due to the high influx of these species from the sewage system (28). *Trichoccocus* was among the most abundant genera observed of the tested anoxic culture conditions, and cultured representatives from this genus are facultative anaerobes that can grow at low temperatures and manifest fermentative growth (29, 30). Similar to *Acidovorax*, multiple studies have identified *Trichococcus* as one of the most prevalent genera in sewage systems (31, 32).

### Process-important bacterial species can be isolated

An important goal of this study was to demonstrate that pure cultures of previously uncultured AS species can be isolated based on our optimized growth conditions. To do this, colonies were randomly picked from plates incubated under oxic conditions containing ASF supplemented with acetate, tryptone, or casamino acids taking care to select both large and small colonies. The colonies were transferred to liquid isolation medium in 96-deep-well plates and incubated for 2 weeks at 25°C under oxic conditions. Hereafter, 16S rRNA gene amplicon sequencing was performed on DNA extracted from 185 growing cultures. Sequencing revealed the enrichment of 102 strains based on unique ASVs. The average read abundance of these ASVs was 67% with 153 out of 185 below 95%, indicating that subsequent single colony isolation was required to obtain pure cultures. Classification of the ASVs using the MiDAS 4 database revealed 48 and 43 distinct genera and species, respectively (Supplementary Data S1). The enriched genera included previously uncultivated genera, such as *Ca*. Brachybacter (OLB8 midas_s_315 based on the MiDAS 4.8.1 taxonomy) (33), AAP99, Ellin6067, and *Ca.* Propionivibrio (*Ca*. Accumulibacter midas_s_29) (22), which are all abundant in WWTPs, and most of the species (29 out of 43) were only classified based on the MiDAS placeholder taxonomy, highlighting that several previously uncultivated taxa can be isolated using the ASF medium.

Performing high-throughput restreaking, colony picking, and microbial profiling without laboratory automation can be resource-intensive. Therefore, a selection of 20 cultures was made based on their significance in wastewater treatment and average abundance in wastewater treatment plants (WWTPs) in Denmark (6). After restreaking and growing these cultures for two weeks on agarose plates with isolation medium, four colonies of each culture were grown in liquid isolation medium. 16S rRNA gene amplicon sequencing on DNA extracted from the liquid cultures showed the successful isolation for 14 of the cultured based on high relative abundance of the ASV of interest (>95%). These liquid cultures along with four additional cultures showing enrichment of interesting strains were used to prepare glycerol stocks for long time storage. The stocks were eventually plated, and the resulting colonies evaluated by amplicon sequencing. This confirmed the successful isolation of 10 species (Table 1 and Supplemental Data S1).

**Table 1.**
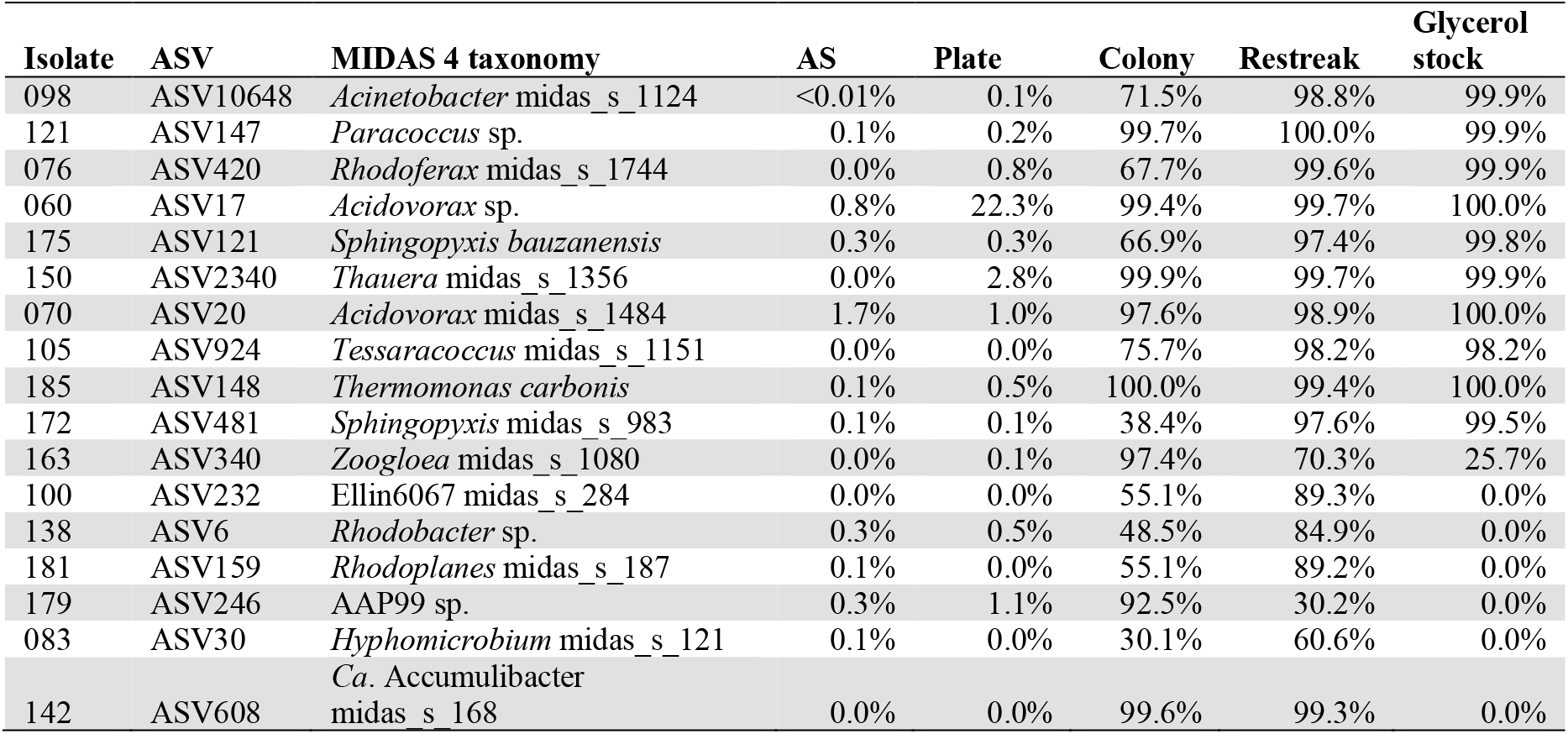
Relative abundance of targeted species throughout the isolation process. AS: ASV abundance in the source activated sludge. Plate: ASV abundance after AS single cells were plated and incubated under oxic conditions. Colony and Restreak: ASV abundance of picked colony from the ASF plate and the subsequent restreak after being incubated in isolation medium for two weeks. Glycerol stock: ASV abundance of colonies after a restreaking the final glycerol stock.

When evaluating the amplicon abundance data for each step of the isolation process several aspects seem to stand out: i) Most species present in high abundance in cultures derived directly from initial culture plates were successfully isolated after several restreaks. Conversely, most species with low abundance for colony forming units did not achieve successful species isolation. ii) Several abundant species on the plates were not able to grow in liquid culture. This could be due to differences in growth conditions since physicochemical conditions might not accommodate the proliferation for the species of interest. Additionally, some species might manifest loss of viability when frozen as glycerol stocks at −80°C during the final step of single colony isolation (34). iii) While it is not unlikely that for some of the species intermediate liquid cultures could have been contaminated by outcompeting bacterial species, sequencing data of for instance the restreaks of *Zoogloea* midas_s_1080 showed consistent presence of both species of interest as well as an *Enterobacter* species (Supplemental Data S1). This seems to indicate that *Zoogloea* midas_s_1080 is not able to grow in isolation, suggesting the presence of an obligate syntrophic relation between the two species (35).

### Most isolates are dependent on ASF for growth

The notable increase in microbial diversity associated with ASF supplementation suggests that most microbial species in AS were dependent on ASF for growth. To determine which components in the ASF promote growth, we plated the 10 isolates on agarose plates with acetate, NH_4_Cl and either no ASF, fresh ASF, autoclaved ASF, or ashed ASF and examined their growth (Figure 4). Autoclaving destroys heat sensitive organic molecules such as signal peptides and certain vitamins in the ASF, whereas ashing removes all organic components, leaving only inorganic compounds, such as minerals and trace elements. We found that none of the isolates were able to grow on acetate and NH_4_Cl alone. However, the addition of fresh ASF promoted the growth of all isolates (Figure 4). Autoclaved ASF was able to recover the growth of seven isolates, whereas not growth was observed with the addition of ashed ASF, indicating an auxotrophy for certain organic metabolites for all the isolates.

**Figure 4.**
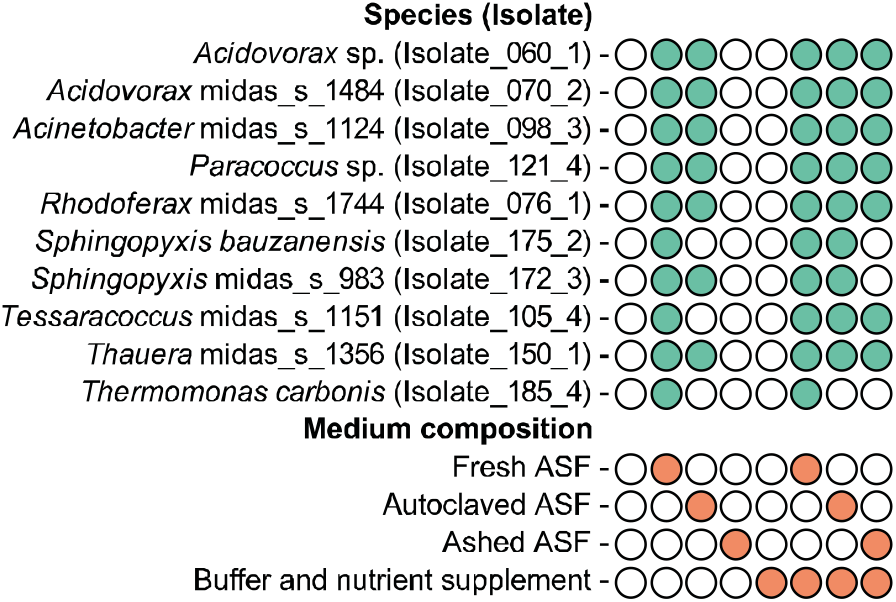
Effect of media composition on the growth of the 10 pure culture isolates. Growth as determined using agarose plates containing 250 mg/L acetate and 50 mg/L NH_4_Cl supplemented with no ASF, fresh ASF, autoclaved ASF, or ashed ASF. Buffer and nutrient supplement (500 mg/L K_2_HPO_4_, 50 mg/L MgSO_4_*7H_2_O, 50 mg/L yeast extract, 300 mg/L sodium pyruvate, 100mg/L casamino acids, and 100 mg/L tryptone) was used to evaluate auxotrophy towards common nutrients. Closed green circles indicate observed growth after two weeks of incubation at 25°C. Closed orange circles indicated addition of the specific medium component.

To determine if growth inhibition was caused by auxotrophy towards common nutrients, we investigated if addition of buffer and nutrient found in R2A growth medium (yeast extract and sodium pyruvate) as well as additional amino acid source (tryptone and casamino acids) could recover the growth of the isolates that were unable to grow on plates lacking ASF or were supplemented with autoclaved or ashed ASF. Interesting, no growth was observed unless some form of ASF was supplemented here as well. This indicates that the ASF provides components other than the common nutrients found in R2A media that are essential for the growth of all 10 species. However, we observed that the addition of the nutrient mix recovered the growth of several isolates that were unable to grow on either autoclaved or ashed ASF. This suggests that most species are also dependent on one or several inorganic compounds that were exclusively present in ASF and could not be reconstituted using the tested nutrient supplement.

To gain more insight into the elemental composition of ASF, ashed ASF, and the nutrient supplement, we analyzed media containing these using ICP/OES (Table 2). The measured concentrations were found to be in the same order of magnitude for most measured elements. One exception was cobalt, which was roughly four times lower in the nutrient supplement when compared to the ASF. The addition of trace elements is known to stimulate microbial conversions and specifically cobalt has been shown to affect overall AS growth (36). It would therefore be interesting to examine the specific effect of cobalt in a future study.

**Table 2.** Concentration of elements (µM) measured for the culture medium containing activated sludge fluid (ASF), ashed ASF, and medium supplement using ICP/OES.

| Element | ASF | Ashed ASF | Nutrient supplement |
| --- | --- | --- | --- |
| Al | 72 | 102 | 61 |
| Ca | 2371 | 1155 | 408 |
| Co | 0.28 | 0.33 | 0.07 |
| Cr | 27 | 59 | 25 |
| Cu | 0.4 | 0.3 | 0.1 |
| Fe | 70 | 179 | 55 |
| Mg | 458 | 201 | 198 |
| Mn | 3.4 | 3.4 | 0.9 |
| Na | 5063 | 5535 | 4142 |
| Ni | 15 | 16 | 5 |
| P | 30 | 1 | 2445 |
| S | 8098 | 584 | 304 |
| Zn | 2 | 1 | 1 |

Aside from the elements measured in our study, it has been shown in several studies that other rare earth metals can be crucial for growth of certain species, including some isolated from extreme habitats (37, 38). While WWTPs influent seems unlikely to be a significant source for these elements (39), further research would be needed to identify any rare earth metal dependencies.

The prior attempt to isolate uncultured bacterial species from bovine rumen proved successful by devising and utilizing a culture medium that faithfully mirrored the mineral composition inherent in the rumen itself (40). Analogously, the growth-promoting effects of ashed ASF might be explicated due to its mineral composition more closely mirroring that within the AS, in comparison to previously employed media like R2A.

### Perspectives

Amongst bacteria that were isolated, successfully cultured and stocked, several novel species are of particular interest for wastewater treatment systems. For instance, species that belong to the *Rhodoferax, Tessaracoccus* and *Thauera* genera have previously been implicated to play a major role in denitrification, phosphate accumulation and floc formation (9). However, isolation of species of some highly desired genera that are present in the AS core community, such as *Ca*. Phosphoribacter (23), *Ca*. Brachybacter (33), and *Ca.* Accumulibacter (22) was unsuccessful. With species abundance data implying that plate culture conditions more closely mimic sewage rather than AS, adjustments could be made by, for example, decreasing concentration of certain substrates further or utilizing plates in which substrates are slowly released (41). Beyond plating-based isolation, the workflow proposed in this study could be combined with alternative cultivation and selection techniques such as dilution to extinction, parallel small scale membrane reactors or cell sorting (4).

Pure cultures of microbes also play a significant role in the transition towards a bio-based economy as, in industry, microorganisms are employed on a massive scale for the production of organic compounds and conversion of resources and as such can provide a plethora of applications in biotechnology (16). However, only several dozen species are commonly used in industry. As with microbial characterization studies, the main obstacle for tapping into the remaining biological resources is our current inability to grow, culture and isolate these species outside of their natural habitat (4).

Incorporating recent developments in lab automation equipment, including pipetting, DNA extraction, plating and colony picking robots into the workflow we present in this study would allow for screening and isolation of bacterial species at substantially higher volumes. As such, the increase in throughput can be used to optimize medium composition specifically for species of particular interest. Combined with PCR based screening methods, this would allow for targeted selection of species that are low in abundance or grow slowly in the selected cultivation conditions.

## Conclusion

This study represents a first exploration of how 16S rRNA gene amplicon sequencing can be used to guide the isolation of novel microbial species directly from AS. ASV-resolved amplicon data classified using the MiDAS 4 reference database provided invaluable information on suitable cultivation conditions for core species in the AS on plates. We found that ASF was required for growth and isolation of several bacterial species known for their high abundance in WWTPs. While initial growth experiments seem to indicate that the ASF mainly substitutes the mineral composition found in WWTPs, further research on the underlying mechanisms would provide valuable insights into what makes a successful cultivation medium.

## Materials and Methods

### Medium preparation

ASF was prepared as follows: 20 L of activated sludge from aeration tanks at AAW WWTP was gathered on 14/10/2020 and 26/01/2021 and ultra-filtrated using Alfa Laval-GR61PP (Alfa Laval, Lund, Sweden) flat disk membranes according to manufacturer specifications (Maximum pressure of 7 bar, cross flow 15 L/min). Subsequently, the permeate was concentrated 2x by reverse osmosis using Alva Laval-R099 (Alfa Laval, Lund, Sweden) flat disk filters using the same operating conditions, and the retentate was filter sterilized using Nalgene^TM^ 0.1 µm bottle-top sterile filter units (Thermo Fisher Scientific, Waltham, MA). For cultivations that required ashed ASF, 500mL of the concentrate was heated to above 100°C to evaporate 95% of the liquid after which the remainder was subjected to 350°C for 2 hours and subsequently reconstituted to 500mL with demineralized water. Autoclaved ASF was obtained by autoclaving the concentrate at 121°C for 20 min.

ASF medium was prepared by supplementing 500 mL of 2x ASF the specified carbon and nitrogen sources to a final concentration of 50 mg/L casamino acids, 100 mg/L mannose, 100 mg/L glucose, 100 mg/L tryptone, 125 mg/L acetate, and 53 mg/L NH_3_Cl using 100x filter-sterilized stock solutions, and adding sterile filter double distilled water (liquid medium) or autoclaved molten 2% DNA pure agarose (VWR, Radnor, PA) (agarose plates) to 1L. Agarose was used instead of agarose and autoclaved alone to reduce the formation of reactive oxygen species which could otherwise inhibit growth sensitive species (19). Tryptic Soy Broth (TSB) (Sigma Aldrich #22092, Burlington, MA) was prepared according to the manufacturer’s instructions. Reasoner’s 2A medium (R2A) was prepared by combining media components in a suitable quantity of demineralized water to accommodate a final concentration of 500 mg/L yeast extract, 300 mg/L sodium pyruvate, 500 mg/L proteose peptone, 500 mg/L glucose, 500 mg/L K_2_HPO_4_, 50 mg/L MgSO_4_ * 7H_2_O and 250 mg/L acetate, then agarose and/or ASF were added after autoclaving at 120°C for 20 min. TSB and R2A agarose plates were solidified using 15 g/L agarose. Single colony isolates were grown and stocked in an isolation medium composed of R2A supplements, except for glucose and proteose peptone, with ASF, 50 mg/L casamino acids, 100 mg/L tryptone, 125 mg/L acetate, and 53 mg/L NH_4_Cl. Growth characterization studies were performed with agarose plate containing a range of compounds with the concentrations as described for the isolation medium.

### Preparation of activated sludge single cells

Activated sludge (30 mL) from aeration tanks was obtained at the AAW WWTP on 16/12/2020 and 03/03/2021 and immediately homogenized in the lab using the RZR 2020 benchtop stirrer (Heidolph, Schwabach, Germany) with glass/Teflon tissue grinder attached (1 min, 2^nd^ gear). Five mL was subsequently sonicated using a Bandelin Sonopuls HD2200 with MS73 probe (Berlin, Germany) set at 60% amplitude with 6x 10 s pulses with 10 s interval. Single cells were separated by centrifugation through a cell strainer with pore sizes of 40 µm (VWR) for 5 min at 3000 ⨉g, followed by centrifugation through a cell strainer with pore sizes of 5 µm (pluriSelect Life Science, Leipzig, Germany) for 2 min at 8600 ⨉g. Cells present in the permeate were counted in a Bürker– Türk counting chamber and subsequently diluted to approx. 10,000 cells/mL.

### Life-dead staining and microscopy

Live/dead staining was performed using the LIVE/DEAD BacLight Bacterial Viability kit (Thermo Fisher Scientific) following manufacturer recommended protocol. Stained cells were analyzed on a white light laser confocal microscope (TCS SP8 X; Leica, Germany).

### Bacterial plating and growth

Agarose plate cultivation was performed at 25°C for at least 2 weeks. Anoxic plate cultivation was done in 3.5 L anaerobic jars with Oxoid™ AnaeroGen™ gas generation sachets (Thermo Fisher Scientific) to remove oxygen to yield levels < 1%. When adequate colony formation was observed, the total combined biomass for each plate was harvested using cell scrapers (VWR) and suspended in 300 µL DNase-free water. Single-species cultures were obtained by picking colonies using 1 µL sterile inoculation loops and transferred to 800 µL liquid isolation medium in 96 deep well 2 mL plates (VWR) which were incubated for at least 2 weeks on a bench top shaker (RT, 200rpm). Intermediate cultures were concentrated by centrifugation to yield 200 µL of which 160 µL was used as input for DNA extraction and 40 µL was mixed with 100 µL of 40% glycerol and stored at −80°C. Final isolated species were grown in liquid isolation medium and stocked by supplementing with glycerol to a final concentration of 30% and stored at −80°C.

### Elemental composition analysis

Elemental analysis of various culture media was performed by using inductively coupled plasma/optical emission spectrometry (ICP/OES), performed on an iCAP 6000 Series (Thermo Scientific, Waltham, MA) as previously described (42).

### DNA extraction

DNA was extracted from 160 µL AS, single cell suspension, suspended biomass from scraped culture plates, or liquid cultures using the FastDNA Spin kit for soil (MP Biomedicals) according to the MiDAS protocol for plate extraction (aau_wwtp_dna_v_8.0) available at https://www.midasfieldguide.org/guide/protocols. DNA concentration and integrity was assessed using a Qubit 3.0 fluorometer (Thermo Fisher Scientific) and an Agilent 2200 Tapestation (Agilent Technologies, CA, USA), respectively.

### 16S rRNA gene amplicon sequencing

V1-V3 amplicons were made using the 27F (5’-AGAGTTTGATCCTGGCTCAG-3’) (43) and 534R (5’-ATTACCGCGGCTGCTGG-3’) (44) primers with barcodes and Illumina adaptors (Integrated DNA Technologies) (45). 25 μL PCR reactions in duplicate were run for each sample using 1X PCRBIO Ultra Mix (PCR Biosystems), 400 nM of both forward and reverse primer, and 10 ng template DNA. PCR conditions were 95°C, for 2 min followed by 20 cycles of 95°C for 20 s, 56°C for 30 s, and 72°C for 60 s, followed by a final elongation at 72°C for 5 min. PCR products were purified using 0.8x CleanNGS beads and eluted in 25 µL nuclease-free water. The 16S rRNA gene V1-V3 amplicon libraries were pooled in equimolar concentrations and paired-end sequenced (2 × 300 bp) on the Illumina MiSeq using v3 chemistry (Illumina, USA). 10 to 20% PhiX control library was added to mitigate low diversity library effects.

16S rRNA gene V1-V3 were processed using usearch v.11.0.667. Amplicon data was processed differently for plates and isolates. For plates, forward and reverse reads were merged using the - fastq_mergepairs command, filtered to remove phiX sequences using usearch -filter_phix, and quality filtered using usearch -fastq_filter with -fastq_maxee 1.0. For, isolates only forward reads were used, and these were filtered to remove phiX sequences using usearch -filter_phix, trimmed to 250 bp using -fastx_truncate -trunclen 250, and quality filtered using usearch -fastq_filter with -fastq_maxee 1.0. All subsequent steps were the same. Dereplication was performed using - fastx_uniques with -sizeout, and amplicon sequence variants (ASVs) were resolved using the usearch -unoise3 command. An ASV-table was created by mapping the quality filtered reads to the ASVs using the usearch -otutab command with the -zotus and -strand plus options. Taxonomy was assigned to ASVs with the MiDAS 4.8.1 database (6) using the usearch -sintax command with -strand both and -sintax_cutoff 0.8 options.

### Amplicon data analyses

Amplicon data was analyzed with R v.4.0.5 (46) through RStudio IDE v.2022.02.3 with the tidyverse v.1.3.1 (https://www.tidyverse.org/), vegan v.2.5 (47), ggplot2 v. 3.3.6 (48), and Ampvis2 v.2.7.9 (49) packages. For alpha diversity analyses, samples were rarefied to 10,000 reads, and alpha diversity (Observed ASVs and inverse Simpsons) was calculated using Ampvis2. Beta diversity distances based on Bray-Curtis (abundance-based) and Jaccard (presence/absence-based) were calculated at the ASV-level using the vegdist function in the vegan R package and visualized by PCoA plots with Ampvis2. Raw data for heatmaps were prepared using Ampvis2 and visualized using ggplot2. Amplicon data for plates were rarefied to 10,000 reads before making the heatmaps to obtain the same sensitivity toward low abundant taxa. Figures were assembled and polished in Adobe Illustrator v.26.3.1.

### Data Availability

The raw sequencing data generated in this study have been deposited in the NCBI SRA database under accession code PRJNA981068.

## Acknowledgement

We thank Emma Abildgaard Thiessen for her help in the lab, Jytte Dencker for conducting the ICP/OES analysis, Zivile Kondrotaite for helping with the confocal microscope and Laura Valk for help with making figures. The project has been funded by the Villum Foundation (Dark Matter and grant 13351, P.H.N.) and the Dutch research council (NWO, employment of MDV).

## Supplementary Information

**Supplementary Information:** Figure S1-S4.

**Supplementary Data S1:** 16S rRNA gene V1-V3 amplicon sequencing results for enrichments and isolates

## References

1. Nielsen PH. 2017. Microbial biotechnology and circular economy in wastewater treatment. Microb Biotechnol 10:1102–1105.

2. Dueholm MKD, Besteman M, Zeuner EJ, Riisgaard-Jensen M, Nielsen ME, Vestergaard SZ, Heidelbach S, Bekker NS, Nielsen PH. 2023. Genetic potential for exopolysaccharide synthesis in activated sludge bacteria uncovered by genome-resolved metagenomics. Water Res 229:119485.

3. Siddharth T, Sridhar P, Vinila V, Tyagi RD. 2021. Environmental applications of microbial extracellular polymeric substance (EPS): A review. J Environ Manage 287:112307.

4. Lewis WH, Tahon G, Geesink P, Sousa DZ, Ettema TJG. 2021. Innovations to culturing the uncultured microbial majority. Nat Rev Microbiol 19:1–16.

5. Wu L, Ning D, Zhang B, Li Y, Zhang P, Shan X, Zhang Q, Brown M, Li Z, Van Nostrand JD, Ling F, Xiao N, Zhang Y, Vierheilig J, Wells GF, Yang Y, Deng Y, Tu Q, Wang A, Zhang T, He Z, Keller J, Nielsen PH, Alvarez PJJ, Criddle CS, Wagner M, Tiedje JM, He Q, Curtis TP, Stahl DA, Alvarez-Cohen L, Rittmann BE, Wen X, Zhou J. 2019. Global diversity and biogeography of bacterial communities in wastewater treatment plants. Nat Microbiol 4:1183–1195.

6. Dueholm MKD, Nierychlo M, Andersen KS, Rudkjøbing V, Knutsson S, Albertsen M, Nielsen PH. 2022. MiDAS 4: A global catalogue of full-length 16S rRNA gene sequences and taxonomy for studies of bacterial communities in wastewater treatment plants. 1. Nat Commun 13:1908.

7. Dueholm MS, Andersen KS, McIlroy SJ, Kristensen JM, Yashiro E, Karst SM, Albertsen M, Nielsen PH. 2020. Generation of comprehensive ecosystem-specific reference databases with species-level resolution by high-throughput full-length 16S rRNA gene sequencing and automated taxonomy assignment (AutoTax). 5. mBio 11:e01557–20.

8. Lobb B, Tremblay BJ-M, Moreno-Hagelsieb G, Doxey AC. 2020. An assessment of genome annotation coverage across the bacterial tree of life. Microb Genomics 6:e000341.

9. Singleton CM, Petriglieri F, Kristensen JM, Kirkegaard RH, Michaelsen TY, Andersen MH, Kondrotaite Z, Karst SM, Dueholm MS, Nielsen PH, Albertsen M. 2021. Connecting structure to function with the recovery of over 1000 high-quality metagenome-assembled genomes from activated sludge using long-read sequencing. 1. Nat Commun 12:2009.

10. Zhang Y, Zhang T. 2022. Culturing the uncultured microbial majority in activated sludge: A critical review. Crit Rev Environ Sci Technol 53:601–624.

11. Nierychlo M, Andersen KS, Xu Y, Green N, Jiang C, Albertsen M, Dueholm MS, Nielsen PH. 2020. MiDAS 3: An ecosystem-specific reference database, taxonomy and knowledge platform for activated sludge and anaerobic digesters reveals species-level microbiome composition of activated sludge. Water Res 182:115955.

12. Heylen K, Lebbe L, De Vos P. 2008. *Acidovorax caeni* sp. nov., a denitrifying species with genetically diverse isolates from activated sludge. Int J Syst Evol Microbiol 58:73–77.

13. Slijkhuis H. 1983. *Microthrix parvicella*, a filamentous bacterium isolated from activated sludge: cultivation in a chemically defined medium. Appl Environ Microbiol 46:832–839.

14. Nowka B, Off S, Daims H, Spieck E. 2015. Improved isolation strategies allowed the phenotypic differentiation of two *Nitrospira* strains from widespread phylogenetic lineages. FEMS Microbiol Ecol 91:fiu031.

15. Shao Y, Chung BS, Lee SS, Park W, Lee S-S, Jeon CO. 2009. *Zoogloea caeni* sp. nov., a floc-forming bacterium isolated from activated sludge. Int J Syst Evol Microbiol 59:526–530.

16. Calero P, Nikel PI. 2019. Chasing bacterial chassis for metabolic engineering: a perspective review from classical to non-traditional microorganisms. Microb Biotechnol 12:98–124.

17. Stewart EJ. 2012. Growing unculturable bacteria. 16. J Bacteriol 194:4151–4160.

18. Yin Q, Sun Y, Li B, Feng Z, Wu G. 2022. The r/K selection theory and its application in biological wastewater treatment processes. Sci Total Environ 824:153836.

19. Tanaka T, Kawasaki K, Daimon S, Kitagawa W, Yamamoto K, Tamaki H, Tanaka M, Nakatsu CH, Kamagata Y. 2014. A hidden pitfall in the preparation of agar media undermines microorganism cultivability. Appl Environ Microbiol 80:7659–7666.

20. Kato S, Yamagishi A, Daimon S, Kawasaki K, Tamaki H, Kitagawa W, Abe A, Tanaka M, Sone T, Asano K, Kamagata Y. e00807-18. Isolation of previously uncultured slow-growing bacteria by using a simple modification in the preparation of agar media. 19. Appl Environ Microbiol 84:1–9.

21. Kämpfer P, Trček J, Skok B, Šorgo A, Glaeser SP. 2015. *Chryseobacterium limigenitum* sp. nov., isolated from dehydrated sludge. Antonie Van Leeuwenhoek 107:1633–1638.

22. Petriglieri F, Singleton CM, Kondrotaite Z, Dueholm MKD, McDaniel EA, McMahon KD, Nielsen PH. 2022. Reevaluation of the phylogenetic diversity and global distribution of the genus “*Candidatus* Accumulibacter.” mSystems 7:e00016–22.

23. Singleton CM, Petriglieri F, Wasmund K, Nierychlo M, Kondrotaite Z, Petersen JF, Peces M, Dueholm MS, Wagner M, Nielsen PH. 2022. The novel genus, ‘*Candidatus* Phosphoribacter’, previously identified as *Tetrasphaera*, is the dominant polyphosphate accumulating lineage in EBPR wastewater treatment plants worldwide. ISME J 16:1605–1616.

24. Joyce E, Al-Hashimi A, Mason T j. 2011. Assessing the effect of different ultrasonic frequencies on bacterial viability using flow cytometry. J Appl Microbiol 110:862–870.

25. Kirkegaard RH, McIlroy SJ, Kristensen JM, Nierychlo M, Karst SM, Dueholm MS, Albertsen M, Nielsen PH. 2017. The impact of immigration on microbial community composition in full-scale anaerobic digesters. Sci Rep 7:9343.

26. Dottorini G, Michaelsen TY, Kucheryavskiy S, Andersen KS, Kristensen JM, Peces M, Wagner DS, Nierychlo M, Nielsen PH. 2021. Mass-immigration determines the assembly of activated sludge microbial communities. Proc Natl Acad Sci U S A 118:e2021589118.

27. Gonzalez-Martinez A, Rodriguez-Sanchez A, Lotti T, Garcia-Ruiz M-J, Osorio F, Gonzalez-Lopez J, van Loosdrecht MCM. 2016. Comparison of bacterial communities of conventional and A-stage activated sludge systems. Sci Rep 6:18786.

28. McIlroy SJ, Kirkegaard RH, McIlroy B, Nierychlo M, Kristensen JM, Karst SM, Albertsen M, Nielsen PH. 2017. MiDAS 2.0: An ecosystem-specific taxonomy and online database for the organisms of wastewater treatment systems expanded for anaerobic digester groups. Database 2017:bax016.

29. Liu J-R, Tanner RS, Schumann P, Weiss N, McKenzie CA, Janssen PH, Seviour EM, Lawson PA, Allen TD, Seviour RJ. 2002. Emended description of the genus *Trichococcus*, description of *Trichococcus collinsii* sp. nov., and reclassification of *Lactosphaera pasteurii* as *Trichococcus pasteurii* comb. nov. and of *Ruminococcus palustris* as *Trichococcus palustris* comb. nov. in the low-G+C Gram-positive bacteria. Int J Syst Evol Microbiol 52:1113–1126.

30. Pikuta EV, Hoover RB, Bej AK, Marsic D, Whitman WB, Krader PE, Tang J. 2006. *Trichococcus patagoniensis* sp. nov., a facultative anaerobe that grows at −5 °C, isolated from penguin guano in Chilean Patagonia. Int J Syst Evol Microbiol 56:2055–2062.

31. VandeWalle JL, Goetz GW, Huse SM, Morrison HG, Sogin ML, Hoffmann RG, Yan K, McLellan SL. 2012. *Acinetobacter*, *Aeromonas*, and *Trichococcus* populations dominate the microbial community within urban sewer infrastructure. Environ Microbiol 14:2538–2552.

32. McLellan SL, Roguet A. 2019. The unexpected habitat in sewer pipes for propagation of microbial communities and their imprint on urban waters. Curr Opin Biotechnol 57:34–41.

33. Kondrotaite Z, Valk LC, Petriglieri F, Singleton C, Nierychlo M, Dueholm MKD, Nielsen PH. 2022. Diversity and ecophysiology of the genus OLB8 and other abundant uncultured *Saprospiraceae* genera in global wastewater treatment systems. Front Microbiol 13:917553.

34. Saheb Alam S, Persson F, Wilén B-M, Hermansson M, Modin O. 2015. Effects of storage on mixed-culture biological electrodes. Sci Rep 5:18433.

35. Connon SA, Giovannoni SJ. 2002. High-throughput methods for culturing microorganisms in very-low-nutrient media yield diverse new marine isolates. Appl Environ Microbiol 68:3878–3885.

36. Gikas P. 2007. Kinetic responses of activated sludge to individual and joint nickel (Ni(II)) and cobalt (Co(II)): An isobolographic approach. J Hazard Mater 143:246–256.

37. Picone N, Op den Camp HJ. 2019. Role of rare earth elements in methanol oxidation. Curr Opin Chem Biol 49:39–44.

38. Smeulders MJ, Barends TRM, Pol A, Scherer A, Zandvoort MH, Udvarhelyi A, Khadem AF, Menzel A, Hermans J, Shoeman RL, Wessels HJCT, van den Heuvel LP, Russ L, Schlichting I, Jetten MSM, Op den Camp HJM. 2011. Evolution of a new enzyme for carbon disulphide conversion by an acidothermophilic archaeon. 7369. Nature 478:412–416.

39. Kawasaki A, Kimura R, Arai S. 1998. Rare earth elements and other trace elements in wastewater treatment sludges. Soil Sci Plant Nutr 44:433–441.

40. Kenters N, Henderson G, Jeyanathan J, Kittelmann S, Janssen PH. 2011. Isolation of previously uncultured rumen bacteria by dilution to extinction using a new liquid culture medium. J Microbiol Methods 84:52–60.

41. Wilming A, Bähr C, Kamerke C, Büchs J. 2014. Fed-batch operation in special microtiter plates: a new method for screening under production conditions. J Ind Microbiol Biotechnol 41:513–525.

42. Jørgensen MK, Nierychlo M, Nielsen AH, Larsen P, Christensen ML, Nielsen PH. 2017. Unified understanding of physico-chemical properties of activated sludge and fouling propensity. Water Res 120:117–132.

43. Usadel B. 1991. 16S/23S rRNA sequencing. In ‘Nucleic acid techniques in bacterial systematics.’ Chichester UK John Wiley Sons 115–175.

44. Parada AE, Needham DM, Fuhrman JA. 2016. Every base matters: assessing small subunit rRNA primers for marine microbiomes with mock communities, time series and global field samples. Environ Microbiol 18:1403–1414.

45. Caporaso JG, Lauber CL, Walters WA, Berg-Lyons D, Huntley J, Fierer N, Owens SM, Betley J, Fraser L, Bauer M, Gormley N, Gilbert JA, Smith G, Knight R. 2012. Ultra-high-throughput microbial community analysis on the Illumina HiSeq and MiSeq platforms. ISME J 6:1621–1624.

46. Team R. 2006. A language and environment for statistical computing. Computing 1.

47. Oksanen J, Blanchet FG, Kindt R, Legendre P, Minchin P, O’Hara B, Simpson G, Solymos P, Stevens H, Wagner H. 2015. Vegan: Community Ecology Package. R Package Version 22-1 2:1–2.

48. Wilkinson L. 2011. ggplot2: Elegant Graphics for Data Analysis by WICKHAM, H. Biometrics 67:678–679.

49. Andersen KS, Kirkegaard RH, Karst SM, Albertsen M. 2018. ampvis2: an R package to analyse and visualise 16S rRNA amplicon data. bioRxiv https://doi.org/10.1101/299537.

