## Supplementary Information for "Amplicon-guided isolation and cultivation of previously uncultured microbial species from activated sludge"

**Affiliations:**

**Table of content:**

Page 2-5: Supplementary Figure 1-4

### Supplementary Figures:

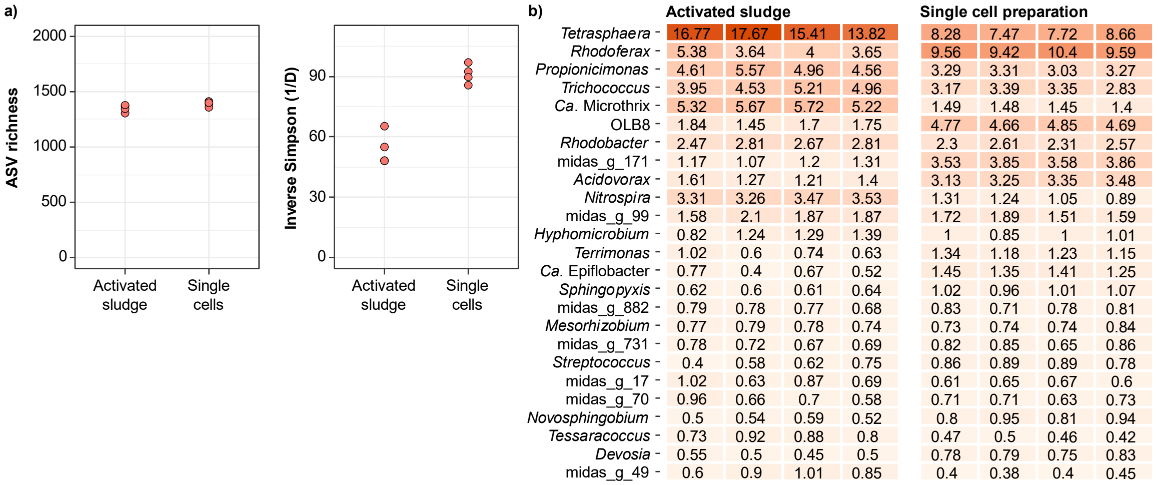


**Figure S1**. Preparation of single cell suspensions from activated sludge. a) Alpha diversity based on 16S rRNA gene V1-V3 amplicon data. b) Heatmap of the 25 most abundant genera in the activated sludge and single cell suspensions. Figures are based on activated sludge collected the 3rd of March 2021 and used for anoxic cultivations.

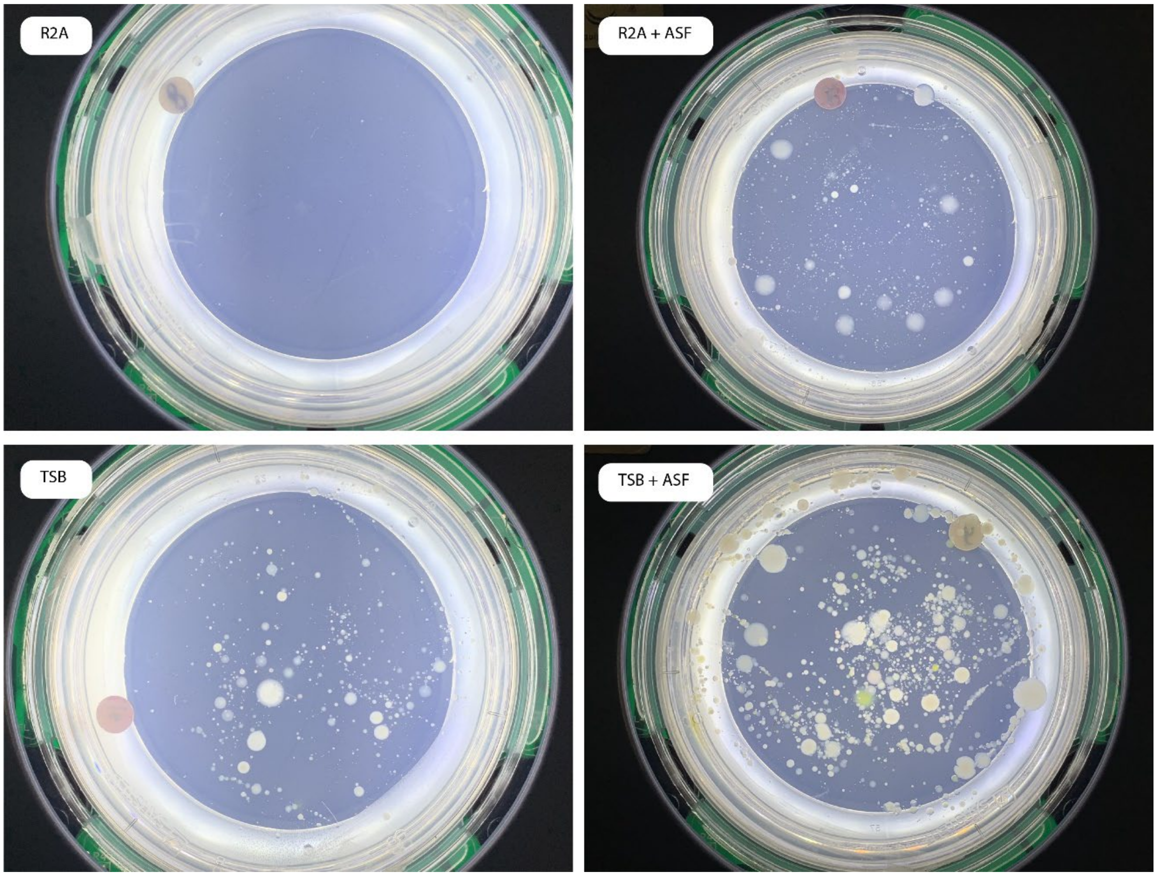


**Figure S2**: Photographs taken from agarose plates with for R2A and TSB medium with and without ASF. For each plate approximately 1000 cells were plated and incubated for 2 weeks at 25°C under oxic conditions.

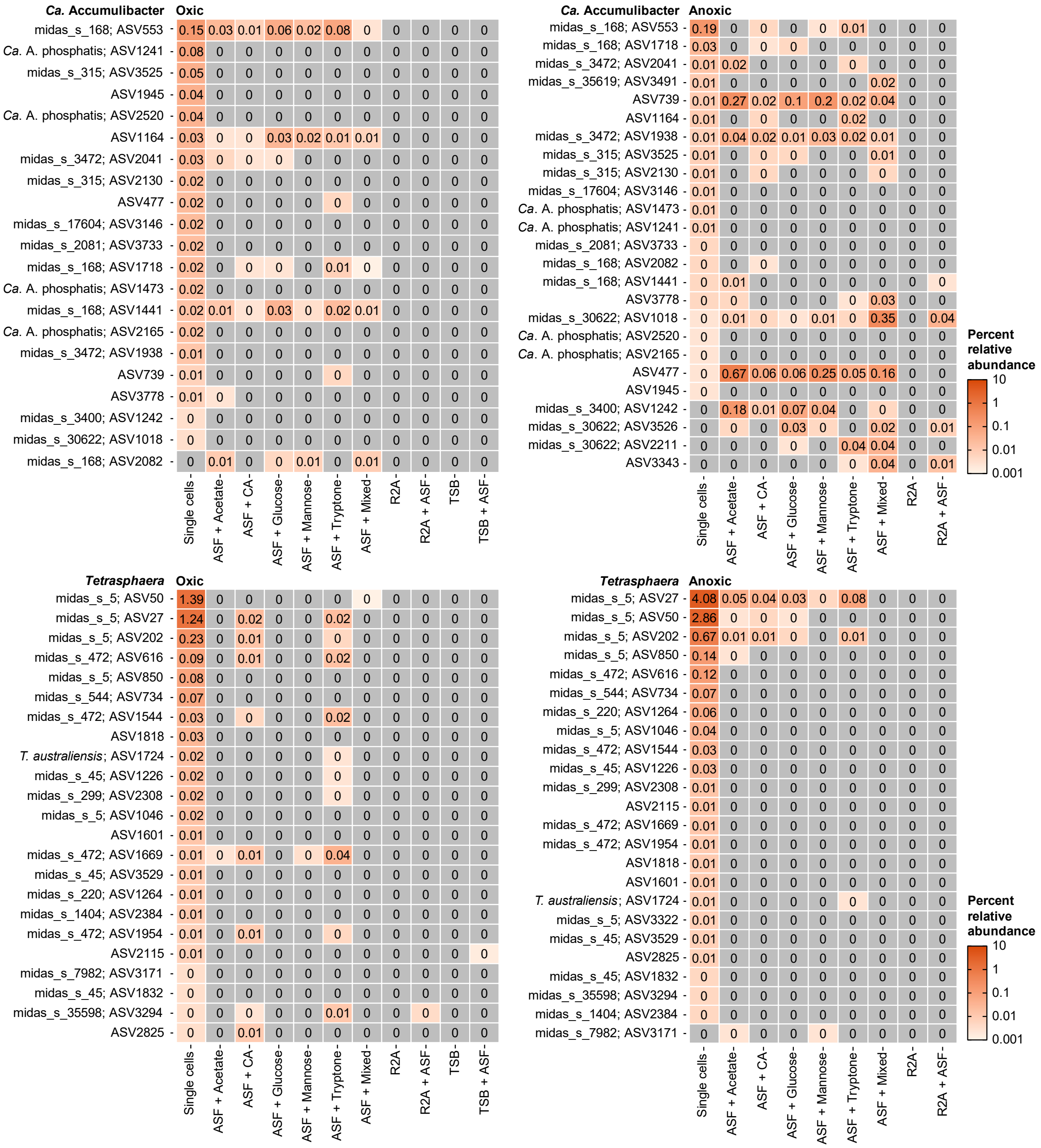


**Figure S3**. Heat map showing all *Ca*. Accumulibacter and *Tetrasphaera* ASVs found in the AS single cell suspension and the corresponding abundancy values found on each plate incubation under oxic and anoxic conditions. Values are the average abundance from four separately processed agarose plates. ASF: Activated sludge fluid; CA: Casamino acids.

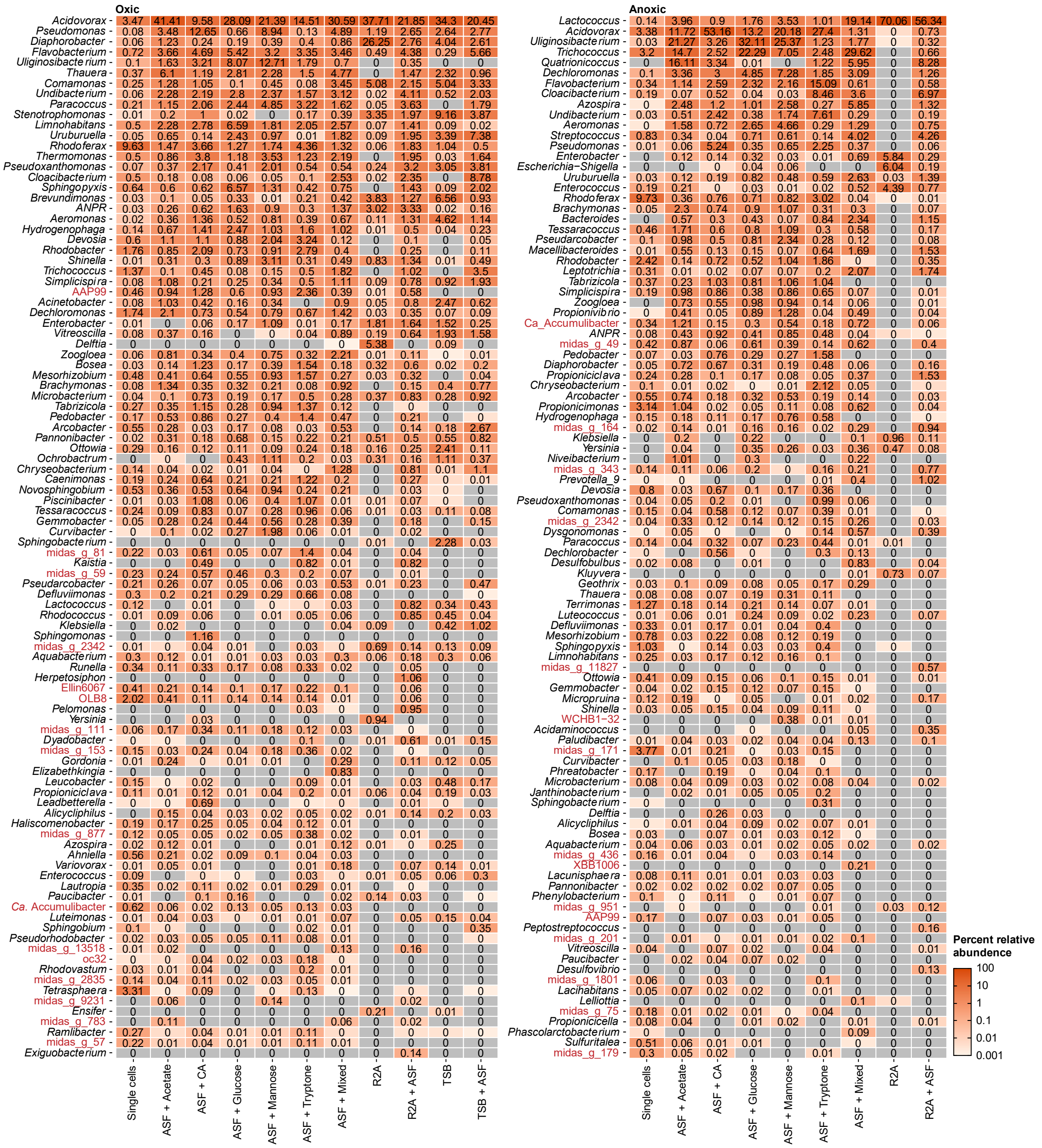


**Figure S4**. Heat map showing the top 100 genera found on the agarose plates after incubation under oxic and anoxic conditions. Values are the average abundance from four separately processed agarose plates, and genera are sorted based on average relative abundance across all media. Genera colored in red have no representative isolates. ASF: Activated sludge fluid; CA: Casamino acids; *ANPR*: *Allorhizobium-Neorhizobium-Pararhizobium- Rhizobium*.
